# Dual Lineage Tracing Identifies Cellular Mechanisms Underlying Radiation-Associated Changes in Atherosclerotic Lesion Composition

**DOI:** 10.64898/2026.01.14.699566

**Authors:** Rebecca A. Deaton, Tajbir Raihan, Victoria M. Milosek, Fatema Allaham, Alexandra L. Krinsky, Laura S. Shankman, Gabriel F. Alencar, Anita Salamon, Nazanin Moradinasab, Subhashis Banerjee, Gary K. Owens

## Abstract

**Background:** Phenotypic plasticity of smooth muscle cells (SMCs) and endothelial cells (ECs) contributes to atherosclerotic plaque composition and stability, yet how shifts in one population influence the contribution and function of the other under conditions of vascular stress, such as irradiation, is poorly understood. A major limitation has been the inability to *simultaneously* fate-map both cell types within the same lesion, with most studies mapping one lineage while inferring the other using unreliable dynamically changing marker genes, risking false-positive and false-negative assignment.

**Methods:** We generated dual lineage-tracing *Apoe*-deficient mice, enabling simultaneous fate mapping of SMCs and ECs. This model was used to extend prior findings from single lineage-tracing models demonstrating irradiation-induced loss of SMC lesion investment and expansion of EC-derived cells. Dual lineage-tracing mice were subjected to irradiation and bone marrow transplantation, followed by Western diet feeding to induce atherosclerosis. Lineage tracing, immunostaining and scRNA-seq analysis were used to define coordinated SMC and EC responses and identify changes relevant to plaque instability.

**Results:** Dual lineage tracing specifically and simultaneously labeled SMC- and EC-derived cells in healthy and atherosclerotic vessels. Irradiation induced divergent responses: SMC-derived cells failed to invest in lesions and upregulated stress-activated inflammatory genes, whereas EC-derived cells expanded and upregulated SMC-associated genes. However, EC-derived cells within lesions failed to induce extracellular matrix genes, and lesions from irradiated mice exhibited reduced collagen content and fewer ACTA2^+^ cells within the fibrous cap, consistent with reduced plaque stability.

**Conclusions:** Dual lineage-tracing of SMCs and ECs demonstrated that irradiation-induced loss of lesional SMC and expansion of EC-derived ACTA2^+^ cells are not artifacts of false lineage assignment. By resolving SMC and EC fate within the same lesion, we identify irradiation-induced cell dynamics including stress-activated inflammatory reprogramming of SMCs, EC phenotypic modulation, impaired extracellular matrix organization, and reduced ACTA2⁺ fibrous cap cellularity that may contribute to radiotherapy-associated increased atherosclerotic cardiovascular disease risk.

**Clinical Perspective:** **What Is New?**

- We developed a dual lineage-tracing mouse model that enables simultaneous fate mapping of smooth muscle cells and endothelial cells within the same atherosclerotic lesion.
- This model reveals coordinated arterial cell wall responses to vascular injury that cannot be resolved using single lineage-tracing approaches.
- Extending prior observations, we show that irradiation-induced inflammatory reprogramming of smooth muscle cells and endothelial-to-mesenchymal transition of endothelial cells towards a smooth muscle cell-like state are associated with reduced total lesion collagen content and decreased overall ACTA2^+^ fibrous cap cellularity.
- This dual lineage-tracing mouse establishes a broadly applicable model for investigating arterial wall cell dynamics across diverse vascular disease states.

**What Are the Clinical Implications?**

- Cancer therapies involving radiotherapy are associated with increased long-term risk of atherosclerotic cardiovascular disease.
- Our findings identify a potential cellular mechanism underlying this risk, in which irradiation-induced smooth muscle cell loss is not functionally compensated by endothelial-to-mesenchymal transition toward a SMC-like state.
- This dual lineage-tracing model provides a tool to evaluate how cancer therapies and other vascular stressors may alter arterial wall cell fate and indices of plaque stability in atherosclerosis and other vascular diseases.

## Introduction

Cardiovascular disease remains the leading cause of death worldwide, with atherosclerotic plaque rupture underlying the morbidity and mortality of most myocardial infarctions and ischemic strokes.^1^ While lipid accumulation and inflammation initiate lesion development, the cellular composition of the plaque ultimately governs its stability through alterations in fibrous cap thickness and extracellular matrix (ECM) deposition and organization.^2,3^ Understanding how distinct vascular cell populations contribute to these processes is therefore central to identifying mechanisms that promote plaque stability.

Lineage tracing studies have fundamentally reshaped our understanding of atherosclerotic plaque biology by revealing the extensive phenotypic plasticity of SMCs and ECs.^4–13^ SMC-specific lineage tracing has shown that SMCs not only contribute to fibrous cap formation but can also give rise to diverse plaque cell phenotypes with context-dependent beneficial or detrimental effects.^5–7,9^ Similarly, EC-specific lineage tracing has demonstrated that ECs contribute to the ACTA2^+^ fibrous cap through EndoMT, challenging the longstanding view that the fibrous cap is derived exclusively from SMCs.^6^ The identification of EC-derived cells in the fibrous cap has implicated ECs in plaque stabilization; however, whether EndoMT promotes plaque stability or vulnerability remains actively debated.^12,14^ Taken together, these studies establish that phenotypic modulation of both SMCs and ECs directly shape plaque composition and stability.

Despite these advances, a major limitation of existing approaches is the reliance on single lineage-tracing models, which preclude simultaneous, rigorous assessment of both SMC- and EC-derived cell populations within the same lesion and often involve mouse lines with different congenic backgrounds, further complicating direct comparisons. Because SMCs and ECs undergo dynamic and potentially interdependent phenotypic changes during atherosclerosis, it remains unclear how alterations in one population influence the fate, function, and compensatory capacity of the other. This gap has hindered mechanistic insight into how coordinated and compensatory SMC and EC responses govern plaque stability in the setting of vascular injury and disease.

This limitation is particularly relevant in the context of radiotherapy which is associated with well-documented long-term increases in atherosclerotic cardiovascular disease risk, especially among cancer survivors.^15–17^ Ionizing radiation induces persistent vascular injury and inflammation, yet the cellular mechanisms linking radiation exposure to plaque destabilization remain poorly defined. Prior studies using separate SMC- or EC-specific lineage-tracing mice demonstrated that irradiation markedly reduces SMC accumulation and increases the contribution of endothelial-derived cells in both the lesion and fibrous cap.^6,18^ However, whether EC phenotypic modulation can functionally compensate for SMC loss, and how SMC and EC contributions are coordinated within the same lesion, cannot be fully addressed using single lineage tracing approaches.

In this study, we addressed this limitation by developing a novel dual lineage-tracing mouse model that enables simultaneous fate mapping of SMCs and ECs in both healthy and atherosclerotic vessels. This model allowed us to extend prior studies from our lab^6,18^ showing coordinated loss of SMC-derived cells and expansion of EC-derived cells following irradiation and bone marrow transfer. We further examined the impact of these shifts in cell composition on ECM production and ACTA2^+^ fibrous cap cellularity, two key indices of plaque stability. Moreover, scRNA-seq analyses provided insight into potential mechanisms underlying these changes, including SMC acquisition of stress-activated inflammatory gene programs and EC acquisition of SMC-marker gene expression. Importantly, this dual lineage-tracing strategy provides a powerful model for dissecting SMC-EC interactions in both healthy and disease states.

## Methods

### Animals

Mice were generated and used in accordance with protocols reviewed and approved by the University of Virginia Animal Care and Use Committees. The *Myh11*-DreER^T2^ mice have been previously described.^9^ The *Cdh5-*CreER^T2^ mouse was generated by Ralf Adams.^19^ The Rosa26*-IR1* Interleaved Reporter mouse was generated by Dr. Bin Zhou.^20^ *Apoe* KO and CD45.1 mice were purchased from Jackson Laboratories (strain #002052 and #002014 respectively). SMC-EC dual lineage-tracing *Apoe* KO mice (herein referred to as SMC-EC dual *Apoe^−/−^*) contain the transgenes necessary to drive SMC-specific Dre recombinase (*Myh11*-DreER^T2^) and EC-specific Cre recombinase (*Cdh5*-CreER^T2^) to recombine the Rosa26-*IR1* reporter resulting in permanent labeling of SMCs and ECs (and their progeny) with tdtomato and zsGreen respectively. Animals were fed standard chow diet (Teklad Irradiated LM-485 Mouse/Rat Diet, Inotiv, 7912) ad libitum. To induce lineage tracing, 6-week-old male and female SMC-EC dual *Apoe^−/−^*mice were fed a tamoxifen diet (250mg tamoxifen/kg diet; Inotiv TD.130856) ad libitum for a period of 12 days before returning to standard chow feed. Tissues from healthy non-atherosclerotic mice were collected for model validation two weeks post tamoxifen diet feeding.

### Irradiation, Bone Marrow Transplant and atherosclerosis

9-week-old male and female SMC-EC dual *Apoe^−/−^* mice underwent lethal irradiation as previously described^6,18^, receiving two 600 cGy doses, 3 hours apart, using a cesium-137 irradiator (Mark 1-68a; JL Shepard and Associates). Bone marrow (BM) was reconstituted 30–60 minutes later with >1 × 10^6^ unfractionated BM cells via tail vein injection. BM was harvested from the femur and tibia from CD45.1 *Apoe^−/−^*donor mice aged 4 to 8 weeks. After BMT, mice were given antibiotics (4.4cc of antibiotic concentrate containing 80mg/mL sulfamethoxazole, 16 mg/mL trimethoprim added to 300mL of drinking water) (Somerset Pharma LLC, NDC 70069-**363**-01) for 6 weeks during BM reconstitution before being placed on WD. Control mice received no irradiation, sterile saline in lieu of donor BM and antibiotic drinking water for 6 weeks to parallel irradiated mice. Following BM reconstitution, control and irradiated mice were placed on Western diet (WD, Inotiv, TD.88137) and fed ad libitum for 18-20 weeks to induce atherosclerosis.

### Statistical Analysis

Statistics were performed using GraphPad Prism 10.6.1. Data are presented as mean ± standard error of the mean (SEM). Normality of data distribution was evaluated using the Shapiro-Wilk test and the *F* test was performed to check for unequal variance. For comparison of two groups of continuous variables with normal distribution and equal variance, two-tailed unpaired Student’s *t*-test (with Welch’s correction when variance was unequal) or for non-normal distribution, two-tailed nonparametric Mann-Whitney *U* test was performed. For datasets involving repeated measures, data were analyzed using repeated measures ANOVA or a mixed-effects model depending on data completeness. Post-hoc comparisons were adjusted using Šidák’s correction when applicable. The number of mice used for each analysis is indicated in the figure legends.

### Data and Code Availability

The data and code supporting the findings of this study are available from the corresponding author upon request.

An expanded methods section can be found in the Supplemental Material.

## Results

### Development and validation of a dual lineage-tracing mouse for coordinated smooth muscle and endothelial cell fate mapping

To enable simultaneous fate mapping of SMCs and ECs withing the same vessel and lesion, we generated a dual recombinase-based lineage-tracing mouse (SMC-EC dual *Apoe*^−/−^) harboring *Myh11*-DreER^T2^ and *Cdh5*-CreER^T2^ transgenes together with the Rosa26-*IR1* dual reporter allele (**Figure 1A**). In this system, tamoxifen-inducible Dre- and Cre-mediated recombination selectively activates tdTomato expression in SMCs and zsGreen expression in ECs respectively.

**Figure 1:**
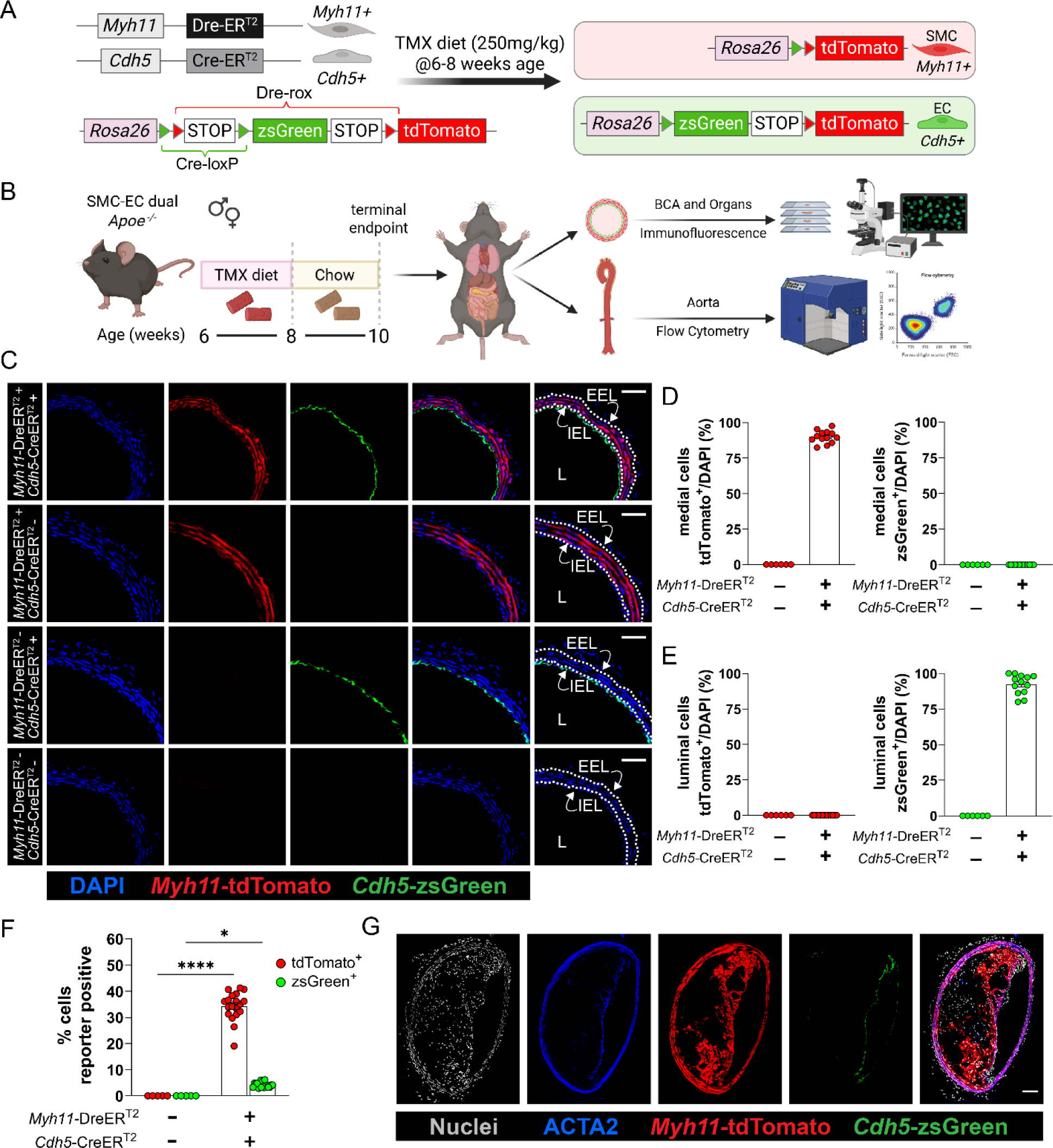
The dual SMC-EC dual *Apoe^−/−^*lineage-tracing mouse labels SMC and EC simultaneously in a cell-type specific manner in healthy and atherosclerotic vessels. (**A**) Schematic describing the SMC-EC dual lineage-tracing mouse model which uses *Myh11*-DreER^T2^ to label SMCs with tdTomato and *Cdh5*-CreER^T2^ to label EC with zsGreen. (**B**) Schematic showing the experimental design for validation studies. (**C**) Representative fluorescent images of cross-sections of brachiocephalic arteries from tamoxifen-fed mice positive (+) for both *Myh11*-DreER^T2^ and *Cdh5*-CreER^T2^ transgenes (top row), positive for only *Myh11*-DreER^T2^ (second row from top), positive for only *Cdh5*-CreER^T2^ (third row from top) or negative (-) for both transgenes (bottom row). (**D**) Quantification of the percent of medial cells (includes all cells between the internal and external elastic lamina [IEL and EEL, respectively] shown in far right panels in C) that are tdTomato^+^ (left) or zsGreen^+^ (right) for mice positive (+) or negative (-) for both the *Myh11*-DreER^T2^ and *Cdh5*-CreER^T2^ transgenes (**E**) Quantification of the percent of luminal cells (cells inside the IEL facing the lumen) that are tdTomato^+^ (left) or zsGreen^+^ (right) for mice positive (+) or negative (-) for both the *Myh11*-DreER^T2^ and *Cdh5*-CreER^T2^ transgenes. (**F**) FACS quantification of the percent of aortic cells that are tdTomato^+^ (red circles) or zsGreen^+^ (green circles) in aortas from mice positive (+) or negative (-) for both the *Myh11*-DreER^T2^ and *Cdh5*-CreER^T2^ transgenes. ****P-value = ˂0.0001, *P-value = 0.0430. (**G**) Representative images of BCA cross-sections from tamoxifen-induced 18 week Western-diet fed SMC-EC dual *Apoe^−/−^*mice. Scale Bar =100µm (**C, G**). n=6 (*Myh11-*DreER^T2^ and *Cdh5-*CreER^T2^ transgene negative [-] mice) and n=13 (*Myh11-*DreER^T2^ and *Cdh5-*CreER^T2^ transgene positive [+] mice) (**D, E**); n=5 (*Myh11-*DreER^T2^ and *Cdh5-*CreER^T2^ transgene negative [-] mice) and n=19 (*Myh11-*DreER^T2^ and *Cdh5-*CreER^T2^ transgene positive [+] mice) (**F**). Bar graphs show the mean ± SEM and biologically independent animals are indicated as individual dots. Statistical significance was assessed using a mixed-effects model with Šidák correction for multiple comparisons (**F**). Schematics were created with BioRender.com

To induce recombinase activity, mice were fed tamoxifen diet for 12 days beginning at 6-8 weeks of age, followed by a two-week washout period prior to tissue collection (**Figure 1B**). Vascular and nonvascular tissues were collected for histological analysis or flow cytometry, and endogenous tdTomato and zsGreen fluorescence was assessed (without antibody amplification) to evaluate cell-type specific labeling.

Representative immunofluorescence images of brachiocephalic artery (BCA) cross-sections demonstrated strict recombinase dependence of reporter activation (**Figure 1C**). Dual lineage labeling required the presence of both recombinase transgenes (**Figure 1C**, top row), whereas mice harboring only one recombinase exhibited labeling of the corresponding cell type alone (*Myh11*-DreER^T2^ only, **Figure 1C**, second row from top; *Cdh5*-CreER^T2^ only, **Figure 1C**, third row from top). Recombinase-negative controls showed no detectable reporter signal (**Figure 1C**, bottom row). Consistent patterns of reporter labeling of vascular and visceral SMCs (tdTomato) and ECs (zsGreen) was observed across multiple tissues in SMC-EC dual *Apoe^−/−^* mice (**Figure S1**).

Quantitative analysis of medial cells in BCAs from SMC-EC dual *Apoe*^−/−^ mice (cells located between the internal elastic lamina [IEL] and the external elastic lamina [EEL] shown in **Figure 1C**, far right panels) showed efficient and specific SMC labeling, with 89.94% ± 1.22% SEM of medial cells expressing tdTomato and no detectable zsGreen^+^ medial cells above the detection limit (see Methods). Conversely, analysis of luminal cells (defined as cells inside the IEL facing the lumen [L] shown in **Figure 1C**, far right panels) showed 92.40% ± 1.92% SEM zsGreen labeling with no detectable tdTomato^+^ luminal cells above the detection limit. As expected, recombinase-negative mice exhibited no detectable reporter labeling. Reporter activation was tamoxifen-dependent, with minimal background recombination in the absence of tamoxifen (**Figure S2**). When stratified by sex, no differences in efficiency or specificity were detected (**Figure S3A-B**), showing consistent lineage tracing in male and female mice.

Flow cytometric analysis of dissociated aortic cell suspensions further confirmed lineage specificity and recombinase dependence (**Figure 1F**). Recombinase-negative mice exhibited negligible reporter-positive cells, whereas SMC-EC dual *Apoe^−/−^* mice displayed both tdTomato^+^ (39.32% ± 1.24% SEM) and zsGreen^+^ (4.26% ± 0.235% SEM) populations. Reporter negative cells reflect additional aortic cell populations, including adventitial fibroblasts and other perivascular tissue as well as any unlabeled SMC and EC. Again, when stratified by sex, no significance differenced were observed by flow cytometry (**Figure S3C**).

Finally, in SMC-EC dual *Apoe*^−/−^ mice fed a Western diet (WD) for 18 weeks to induce the formation of advanced atherosclerotic lesions, reporter labeling not only marked medial SMC and luminal ECs but also identified tdTomato^+^ SMC-derived cells and zsGreen^+^ EC-derived cells within the lesion, consistent with previous studies using the single SMC or EC lineage-tracing mice. Thus, the SMC-EC dual *Apoe*^−/−^lineage-tracing mouse provides a model to investigate alterations in plaque cellular composition in atherosclerosis.

### Irradiation induced coordinated shifts in smooth muscle and endothelial cell contributions associated with reduced indices of plaque stability

To extend our prior studies using separate SMC- and EC-specific lineage-tracing models^6,18^, SMC-EC dual *Apoe^−/−^* mice were subjected to irradiation and bone marrow transplantation followed by WD feeding to induce advanced atherosclerosis (**Figure 2A**). Efficient bone marrow engraftment was confirmed in all irradiated mice (**Figure S4**).

**Figure 2:**
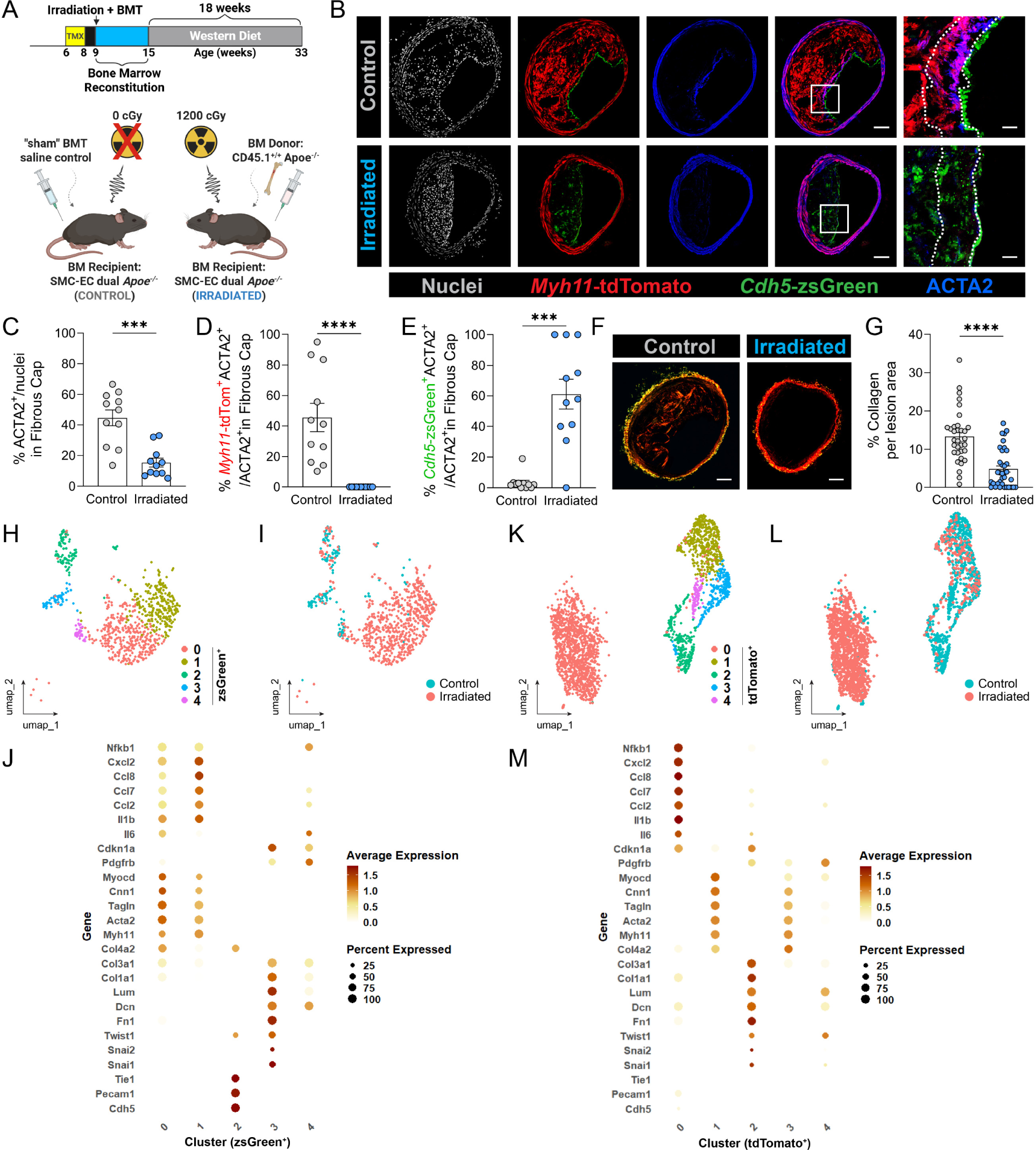
Irradiation drives coordinated but divergent smooth muscle and endothelial cell responses in the same atherosclerotic lesion, accompanied by reduced indices plaque stability. (**A**) Schematic showing the experimental design for irradiation and atherosclerosis studies. (**B**) Representative immunofluorescent images of brachiocephalic artery (BCA) lesion cross-sections from control (top panels) or irradiated (bottom panels) SMC-EC dual *Apoe^−/−^* mice. Panels 1-5 from left to right show nuclei (panel 1), ACTA2 (panel 2), tdTomato and zsGreen (panel 3), merged image (panel 4) and a zoomed image of the fibrous cap (white square in panel 4; zoom shown in panel 5). (**C**) Quantification of ACTA2^+^ cells in the 30μm fibrous cap of BCA atherosclerotic lesions from control (gray circles) or irradiated (blue circles) SMC-EC dual *Apoe*^−/−^ mice. ***P-value = 0.0003. (**D**) Quantification of the percent of total ACTA2^+^ cells that are derived from SMCs (tdTomato^+^ ACTA2^+^) in the 30μm fibrous cap of BCA atherosclerotic lesions from control (gray circles) or irradiated (blue circles) SMC-EC dual *Apoe*^−/−^ mice. ****P-value = ˂0.0001. (**E**) Quantification of the percent of total ACTA2^+^ cells that are derived from ECs (zsGreen^+^ ACTA2^+^) in the 30μm fibrous cap of BCA atherosclerotic lesions from control (gray circles) or irradiated (blue circles) SMC-EC dual *Apoe*^−/−^ mice. ***P-value = 0.0003. (**F**) Representative Picrosirius Red stained BCA atherosclerotic lesions from control (left) or irradiated (right) SMC-EC dual *Apoe*^−/−^ mice. (**G**) Quantification of the total collagen content in the BCA atherosclerotic lesions from control (gray circles) or irradiated (blue circles) SMC-EC dual *Apoe*^−/−^ mice. ****P-value = ˂0.0001. (**H-I**) UMAP of ECs expressing zsGreen transcript, identifying five transcriptionally distinct subclusters (clusters 0–4). Cells are colored by subcluster identity (**H**) or experimental condition (control vs. irradiated; **I**). Dot plot (**J**) shows expression of selected genes across zsGreen⁺ subclusters, where dot size represents the percentage of cells expressing each gene and color intensity indicates average scaled expression. (**K-L**) UMAP of SMCs expressing tdTomato transcript, identifying five transcriptionally distinct subclusters (clusters 0–4). Cells are colored by subcluster identity (**K**) or experimental condition (control vs. irradiated; **L**). Dot plot (**M**) shows expression of selected genes across tdTomato⁺ subclusters, where dot size represents the percentage of cells expressing each gene and color intensity indicates average scaled expression. Scale Bar = 100µm (**B**, panels 1-4 from left) or 20 µm (**B**, panel 5 from left). Bar graphs show the mean ± SEM and biologically independent animals are indicated as individual dots (**C, D, E, F**). Sample sizes were n = 11 (control) and n = 11 (irradiated) **(C-E)**; n = 33 (control) and 35 (irradiated) (**G**). Statistical significance was assessed by two-tailed nonparametric Mann-Whitney *U* test (**C-E, G**). Schematics were created with BioRender.com

Consistent with prior reports demonstrating vascular bed-specific reductions in lesion size following irradiation (specifically the aortic arch and thoracic aorta)^21,22^, we observed reduced lesion area in the BCA. Representative images of MOVAT stained BCA sections are shown for WD-fed control and irradiated mice (**Figure S5A**). BCAs from irradiated mice had diminished outward remodeling, as evidenced by reduced EEL and IEL areas, without significant change in medial area (**Figure S5B-D**). Although both lesion area and percent occlusion were decreased, lumen area remained smaller, reflecting reduced compensatory expansion rather than increased relative plaque burden (**Figure S5E-G**). No sex-specific differences were detected within treatment groups following stratification by sex (**Figure S6**). Because plaque size alone does not fully reflect lesion vulnerability, we next examined changes in cellular composition and indices of lesion stability.

Immunofluorescence analysis of BCA lesions revealed marked irradiation-induced changes in plaque cellular composition that could be readily resolved within the same lesion using dual lineage tracing (representative images shown in **Figure 2B**). Lesions from non-irradiated control mice contained abundant SMC-derived cells (**Figure 2B**, top row, second panel from left). In contrast, lesions from irradiated mice were devoid of SMC-derived cells, shown by the lack of tdTomato^+^ cells and were instead comprised of EC-derived cells, shown by the presence of zsGreen^+^ cells in the lesions (**Figure 2B**, bottom row, second panel from left). Moreover, lesions from irradiated mice had significantly less ACTA2^+^ cells in the 30μm fibrous cap region (**Figure 2B**, bottom row, third and fifth panels from left) compared to control mice (**Figure 2B**, top row, third and fifth panels from left). Quantification of fibrous cap cellularity demonstrated a >65% reduction in total ACTA2^+^ cells in the fibrous cap (from 44.68% ±5.29% SEM to 15.38% ± 2.91% SEM) in lesions from irradiated mice compared to control mice (**Figure 2C**). Consistent with impaired SMC investment in the lesion, the proportion of tdTomato^+^ ACTA2^+^ cells contributing to the ACTA2^+^ fibrous cap was reduced from 45.58% ± 9.28% SEM to 0% ± 0% SEM (no detectable tdTomato^+^ ACTA2^+^ cells) in lesions from irradiated mice compared to control mice (**Figure 2D**). Conversely, the proportion of zsGreen^+^ ACTA2^+^ cells contributing to the ACTA2^+^ fibrous cap increased from 3.29% ± 1.62% SEM to 61.08% ± 9.71% SEM in lesions from irradiated mice compared to control mice (**Figure 2E**) indicating an expansion of EC-derived cells expressing ACTA2. No sex-specific differences in cellular plaque composition were detected within treatment groups following stratification by sex (**Figure S7**).

Phenotypically modulated SMCs and ECs have been reported to express ECM genes^6,9,12^, but whether they deposit and organize ECM proteins to a comparable extent remains unclear. To determine whether coordinated loss of SMC-derived cells and expansion of EC-derived cells in lesions from irradiated mice were accompanied by changes in ECM deposition, picrosirius red (PSR) staining was performed. Representative PSR-stained BCA sections are shown for control and irradiated mice (**Figure 2F**). Quantitative analysis of PSR staining showed a >60% reduction in total collagen content (from 13.32% ± 1.21% SEM to 4.82% ± 0.87% SEM; expressed as a percentage of lesion area) in lesions from irradiated mice compared to control mice (**Figure 2G**). No sex-specific differences in the percent total collagen were detected within treatment groups following stratification by sex (**Figure S8**). These findings indicate that the irradiation-associated shift in cellular composition is accompanied by impaired collagen deposition. Taken together with the reduction in ACTA2^+^ cells within the fibrous cap, these data suggest that lesions from irradiated mice exhibit reduced indices of plaque stability compared with non-irradiated controls.

To gain mechanistic insight into the cellular and transcriptional programs underlying irradiation-induced shifts in cellular contribution and altered ECM deposition, we performed single cell RNA sequencing (scRNA-seq) of cells isolated from the BCA regions of WD-fed control and irradiated SMC-EC dual *Apoe*^−/−^ mice. BCA regions from four control and three irradiated mice were enzymatically dissociated, pooled by treatment group, and subjected to scRNA-seq analysis, generating one dataset per experimental condition. (**Figure S9A**). This approach captures the aggregate transcriptional profile across animals within each group but precludes assessment of biological variance between individual mice.

This approach yielded 27,085 cells in the control dataset and 32,420 cells in the irradiated dataset. Among these, 2,484 control cells (9.17%) and 2,660 irradiated cells (8.20%) expressed *tdTomato* transcript, while 732 control cells (2.70%) and 1,249 irradiated cells (3.85%) expressed *zsGreen* transcript. These differences correspond to an approximately 11% relative reduction in tdTomato^+^ cells and an ∼43% relative increase in zsGreen^+^ cells in irradiated compared with control datasets. These shifts mirror our histologic observations of reduced SMC investment and increased EC contribution in irradiated lesions.

Consistent with prior studies from our laboratory and others^6,9,11,13^, Principal Component Analysis (PCA) followed by graph-based clustering using the Louvain algorithm identified 17 distinct transcriptomic clusters representing the major cell types found in blood vessels and atherosclerotic plaques (**Figure S9B**). Clusters were classified based on canonical marker gene expression (**Figure S9C**). Visualization of cells by experimental treatment (control versus irradiated dataset) revealed a discrete cluster (cluster 0) composed almost exclusively of cells from the irradiated dataset (**Figure S9D**), suggesting that irradiation promoted the emergence of a transcriptionally distinct cell state. Notably, this irradiation-associated cluster contained cells expressing both tdTomato and zsGreen transcripts, indicating contributions from both SMC and EC lineages (**Figure S9E-F**).

Given the enrichment of lineage traced ECs within the irradiation-associated cluster together with our fluorescence imaging analysis demonstrating prominent EC investment in atherosclerotic lesions from irradiated mice, we performed focused subclustering of zsGreen transcript-positive cells. Subclustering revealed five transcriptomically distinct EC-derived subpopulations of EC-derived cells (EC subclusters 0-4, **Figure 2H**). Visualization by treatment group showed that zsGreen^+^ subclusters 0 and 1 were derived almost exclusively from the irradiated dataset, whereas zsGreen^+^ subclusters 2, 3, and 4 contained cells from both the control and irradiated datasets (**Figure 2I**), indicating selective enrichment of specific EC-derived cell populations following irradiation.

Analysis of marker gene expression across the five zsGreen^+^ subclusters highlighted marked phenotypic heterogeneity (**Figure 2J**). zsGreen^+^ subcluster 2 exhibited a canonical endothelial transcriptional profile, with high expression of *Cdh5, Pecam1,*and *Tie1*. In contrast, zsGreen^+^ subclusters 3 and 4 displayed features consistent with EndoMT, characterized by the loss of EC markers and the acquisition of ECM-associated gene expression. zsGreen^+^ subcluster 3 demonstrated a pronounced mesenchymal-like program, with elevated expression of EndoMT-associated transcription factors *Snai1, Snai2* (*Slug*), *Twist1*, *Fn1, Dcn, Lum, Col1a1,* and *Col3a1*, consistent with an advanced EndoMT state. zsGreen^+^ subcluster 4 exhibited a similar but attenuated gene expression profile, suggestive of a partial EndoMT phenotype.

In contrast to these EndoMT-like populations, the irradiation-induced zsGreen^+^ subclusters 0 and 1 showed minimal induction of ECM genes despite loss of canonical endothelial markers. Instead, these subclusters were distinguished by expression of smooth muscle-associated genes including *Myh11, Acta2, Tagln, Cnn1,* and *Myocd*, indicative of a shift toward a SMC-like transcriptional program, accompanied by moderate induction of inflammatory genes (*Il6, Il1b, Ccl2, Ccl7, Ccl8, Cxcl2,* and *Nfkb1*). Together, these findings suggest that irradiation promotes the emergence of EC-derived cells that preferentially acquire SMC-like features rather than undergoing classical EndoMT. This switch in phenotypic modulation, characterized by limited ECM gene expression may contribute to the reduced collagen content observed in the atherosclerotic lesions from irradiated mice.

To further define irradiation-associated changes within the SMC-lineage, we next performed focused subclustering of tdTomato transcript-positive cells, which identified five transcriptionally distinct subpopulations (tdTomato^+^ subclusters 0-4; **Figure 2K**). When visualized by treatment group, tdTomato^+^ subcluster 0 was derived almost exclusively from the irradiated dataset (**Figure 2L**). In contrast, tdTomato^+^ subclusters 1 and 3 contained cells from both control and irradiated datasets, whereas tdTomato^+^ subclusters 2 and 4 were predominantly composed of cells from the control dataset.

Examination of marker gene expression revealed substantial heterogeneity among SMC-derived subclusters (**Figure 2M**). tdTomato^+^ subcluster 1 exhibited a canonical SMC transcriptional profile, characterized by expression of classical SMC marker genes including *Myh11, Acta2, Tagln,* and *Cnn1*. These markers were also expressed by tdTomato^+^ subcluster 3; however, this subcluster showed reduced *Myocd* expression and increased *Pdgfrb*, consistent with an early phenotypically modulated SMC state. tdTomato^+^ subcluster 4 displayed marked downregulation of canonical SMC genes alongside upregulation of *Pdgfrb* and ECM genes including *Lum* and *Dcn* indicative of a transition towards a myofibroblast-like state. tdTomato^+^ subcluster 2 exhibited features consistent with a myofibroblast-like or fibrotic-like SMC phenotype commonly observed in advanced atherosclerotic lesions, with high expression of *Fn1, Lum, Dcn, Col1a1,* and *Col3a1*.

In contrast to these previously described SMC phenotypic states, irradiation-associated tdTomato^+^ subcluster 0 exhibited loss of canonical SMC marker gene expression with minimal induction of ECM genes. Instead, cells within this subcluster were distinguished by pronounced upregulation of stress- and inflammation-associated genes commonly linked to senescence-associated programs, including *Cdkn1a* (*p21*), *Il6, Il1b, Ccl2, Ccl7, Ccl8, Cxcl2* and *Nfkb1*. This transcriptional profile is consistent with an alternative mode of SMC phenotypic modulation characterized by a stress-activated inflammatory state, potentially senescence-like in nature, rather than acquisition of a matrix-producing, migratory or proliferative phenotype. Notably, platelet-derived growth factor receptor β (*Pdgfrb*), which is required for SMC investment into atherosclerotic lesions^6^, was not detected in tdTomato^+^ subcluster 0 cells. Together, these data support a model in which irradiation induces a stress-activated inflammatory program in SMCs, with failure to upregulate *Pdgfrb* expression that may limit their responsiveness to migratory cues and contribute to reduced SMC investment into the lesions of irradiated mice.

## Discussion

Radiation exposure is a well-established risk factor for accelerated cardiovascular disease in cancer survivors^15–17^. However, the cellular mechanisms linking irradiation to altered plaque biology remain incompletely understood. In this study, we leveraged a novel dual SMC-EC lineage-tracing mouse model in combination with histological, molecular and single-cell transcriptomic analysis to define how irradiation reshapes vascular cell fate and function within atherosclerotic lesions. This integrated approach enabled direct tracking of both SMC- and EC-derived cells and revealed irradiation-induced cellular states that would not be discernible using single lineage-tracing models or conventional marker-based approaches alone.

Despite an overall reduction in lesion size and percent occlusion in irradiated mice, plaques exhibited altered outward remodeling, reduced percent of ACTA2^+^ fibrous cap cells and diminished collagen deposition which are all features associated with reduced plaque stability. These findings highlight that lesion burden alone is insufficient to capture the vascular consequences of irradiation and underscore the importance of assessing cellular composition and function. Using SMC-EC dual lineage tracing, we observed a marked shift in plaque cellular investment, with reduced SMC-derived contributions and increased accumulation of EC-derived cells within lesions from irradiated mice.

Single-cell RNA sequencing further demonstrated the utility of this dual lineage tracing strategy. Global clustering identified a transcriptionally distinct cell population almost exclusively associated with irradiation treatment that contained lineage-traced cells from both SMC and EC origins. By integrating lineage information with transcriptomic profiling, we were able to disentangle phenotypically convergent cell states and reveal that irradiation promotes unexpected reprogramming trajectories within both SMC and EC lineages. Notably, focused analysis of EC-lineage cells showed that irradiation preferentially enriched EC-derived populations that lost canonical endothelial identity and adopted SMC-like transcriptional programs, rather than undergoing classical endothelial-to-mesenchymal transition (EndoMT) characterized by extracellular matrix production.

In parallel, subclustering of SMC-derived cells identified a distinct irradiation-associated SMC state defined by loss of contractile genes, minimal induction of ECM, and activation of stress- and inflammation-associated pathways. Cells within this tdTomato^+^ subcluster exhibited elevated levels of *Cdkn1a* (*p21*) together with pro-inflammatory cytokines and chemokines commonly associated with senescence-related and inflammatory states, while lacking expression of *Pdgfrb*, a receptor required for SMC investment within atherosclerotic lesions.^18^ This transcriptional profile parallels prior observations of radiation-induced senescence-associated β-galactosidase activity in medial vascular SMCs, supporting the possibility that irradiation engages senescence-related programs within this SMC subpopulation.^23^ Notably, BCA SMCs are neural crest-derived, and other neural crest-derived lineages, including melanocytes, have been shown to undergo ionizing radiation-induced senescence, providing additional biological support for the senescence-associated transcriptional features observed here.^24^ Together, these findings suggest that irradiation diverts SMCs away from reparative, lesion-stabilizing phenotypes toward a state with limited capacity for migration, proliferation, and matrix deposition, thereby impairing SMC contribution to lesion stabilization. Future studies incorporating functional assays will be required to determine whether this irradiation-associated SMC state represents bona fide cellular senescence or a related stress-induced inflammatory phenotype.

Together, these data support a model in which irradiation increases cardiovascular risk not by increasing plaque size, but by fundamentally altering SMC and EC phenotypic modulations in ways that compromise plaque stability. The emergence of stress-activated inflammatory SMCs and the accumulation of EC-derived cells adopting non-canonical identities may limit effective fibrous cap formation and collagen deposition, providing a potential cellular mechanism linking radiation exposure to adverse plaque composition despite modest plaque burden. This dual SMC-EC lineage-tracing model provides the scientific community with a powerful resource for resolving SMC and EC origin and fate in complex disease settings characterized by extensive phenotypic reprogramming. Future studies leveraging this model may enable the identification of therapeutic strategies aimed at preserving reparative vascular cell states in radiation-exposed populations. Beyond irradiation-associated vascular disease, this model will be broadly applicable to the study of vascular remodeling in other contexts, including aging, metabolic disease, and therapeutic interventions that impact cardiovascular risk.

## Acknowledgements

The authors acknowledge Moon Snyder and Alison Rinehart for their histology expertise, Rupa Tripathi for preparation of reagents required for enzymatic dissociation of cells for scRNA-sequencing, and Jeremy Gatesman for assistance with tail vein injection. The authors also acknowledge the University of Virginia Advanced Microscopy Core (RRID:SCR_018736), the University of Virginia Flow Cytometry Core (RRID:SCR_017829), the Genome Analysis and Technology Core (RRID:SCR_018883), the Bioinformatics Core (RRID:SCR_012718), and the University of Virgina Environment Health and Safety - Radiation Safety team for their assistance in completing the studies described in this manuscript. Pilot analyses were performed using the LECL nuclei detection/classification pipeline for tdTomato and zsGreen; these results were used for internal comparison and were not included in the reported quantification.^25^

## Supplemental Materials List

Supplemental Methods

Figures S1-S9

References

## Nonstandard Abbreviations and Acronyms

ACTA2: smooth muscle alpha actin
Apoe: apolipoprotein E gene
BM: bone marrow
BMT: bone marrow transplant
BCA: brachiocephalic artery
cGy: centigray
EC: endothelial cell
ECM: extracellular matrix
EndoMT: Endothelial-to-mesenchymal transition
FMO: Fluorescence Minus One
scRNA-seq: single-cell RNA sequencing
SMC: smooth muscle cell
UMAP: Uniform Manifold Approximation and Projection
WD: Western diet

## References

1. Martin SS, Aday AW, Allen NB, Almarzooq ZI, Anderson CAM, Arora P, Avery CL, Baker-Smith CM, Bansal N, Beaton AZ, et al. 2025 Heart Disease and Stroke Statistics: A Report of US and Global Data From the American Heart Association. Circulation. 2025;151:e41–e660.

2. Virmani R, Kolodgie FD, Burke AP, Farb A, Schwartz SM. Lessons From Sudden Coronary Death. Arterioscler. Thromb. Vasc. Biol. 2000;20:1262–1275.

3. Bentzon JF, Otsuka F, Virmani R, Falk E. Mechanisms of Plaque Formation and Rupture. Circ. Res. 2014;114:1852–1866.

4. Karnewar S, Karnewar V, Deaton RA, Shankman LS, Benavente ED, Williams CM, Bradley X, Alencar GF, Bulut GB, Kirmani S, et al. IL-1β Inhibition Partially Negates the Beneficial Effects of Diet-Induced Atherosclerosis Regression in Mice. Arterioscler. Thromb. Vasc. Biol. 2024;44:1379–1392.

5. Shankman LS, Gomez D, Cherepanova OA, Salmon M, Alencar GF, Haskins RM, Swiatlowska P, Newman AAC, Greene ES, Straub AC, et al. KLF4-dependent phenotypic modulation of smooth muscle cells has a key role in atherosclerotic plaque pathogenesis. Nat. Med. 2015;21:628–637.

6. Newman AA, Serbulea V, Baylis RA, Shankman LS, Bradley X, Alencar GF, Owsiany K, Deaton RA, Karnewar S, Shamsuzzaman S, et al. Multiple cell types contribute to the atherosclerotic lesion fibrous cap by PDGFRβ and bioenergetic mechanisms. Nat. Metab. 2021;3:166–181.

7. Cherepanova OA, Gomez D, Shankman LS, Swiatlowska P, Williams J, Sarmento OF, Alencar GF, Hess DL, Bevard MH, Greene ES, et al. Activation of the pluripotency factor OCT4 in smooth muscle cells is atheroprotective. Nat. Med. 2016;22:657–665.

8. Gomez D, Baylis RA, Durgin BG, Newman AAC, Alencar GF, Mahan S, St. Hilaire C, Müller W, Waisman A, Francis SE, et al. Interleukin-1β has atheroprotective effects in advanced atherosclerotic lesions of mice. Nat. Med. 2018;24:1418–1429.

9. Alencar GF, Owsiany KM, Karnewar S, Sukhavasi K, Mocci G, Nguyen AT, Williams CM, Shamsuzzaman S, Mokry M, Henderson CA, et al. Stem Cell Pluripotency Genes Klf4 and Oct4 Regulate Complex SMC Phenotypic Changes Critical in Late-Stage Atherosclerotic Lesion Pathogenesis. Circulation. 2020;142:2045–2059.

10. Serbulea V, Martin JM, Owens GK. Role of Diverse Smooth Muscle Cell Phenotypic Transitions in Atherosclerosis Development and Late-Stage Pathogenesis. Annu. Rev. Physiol. 2025.

11. Wirka RC, Wagh D, Paik DT, Pjanic M, Nguyen T, Miller CL, Kundu R, Nagao M, Coller J, Koyano TK, et al. Atheroprotective roles of smooth muscle cell phenotypic modulation and the TCF21 disease gene as revealed by single-cell analysis. Nat. Med. 2019;25:1280–1289.

12. Gole S, Tkachenko S, Masannat T, Baylis RA, Cherepanova OA, Gole S, Tkachenko S, Masannat T, Baylis RA, Cherepanova OA. Endothelial-to-Mesenchymal Transition in Atherosclerosis: Friend or Foe? Cells. 2022;11.

13. Chappell J, Harman JL, Narasimhan VM, Yu H, Foote K, Simons BD, Bennett MR, Jørgensen HF. Extensive Proliferation of a Subset of Differentiated, yet Plastic, Medial Vascular Smooth Muscle Cells Contributes to Neointimal Formation in Mouse Injury and Atherosclerosis Models. Circ. Res. 2016;119:1313–1323.

14. Huang Q, Gan Y, Yu Z, Wu H, Zhong Z. Endothelial to Mesenchymal Transition: An Insight in Atherosclerosis. Front. Cardiovasc. Med. 2021;8:734550.

15. Sturgeon KM, Deng L, Bluethmann SM, Zhou S, Trifiletti DM, Jiang C, Kelly SP, Zaorsky NG. A population-based study of cardiovascular disease mortality risk in US cancer patients. Eur. Heart J. 2019;40:3889–3897.

16. van Nimwegen FA, Schaapveld M, Cutter DJ, Janus CPM, Krol ADG, Hauptmann M, Kooijman K, Roesink J, van der Maazen R, Darby SC, et al. Radiation Dose-Response Relationship for Risk of Coronary Heart Disease in Survivors of Hodgkin Lymphoma. J. Clin. Oncol. 2016;34:235–243.

17. Florido R, Daya NR, Ndumele CE, Koton S, Russell SD, Prizment A, Blumenthal RS, Matsushita K, Mok Y, Felix AS, et al. Cardiovascular Disease Risk Among Cancer Survivors: The Atherosclerosis Risk In Communities (ARIC) Study. J. Am. Coll. Cardiol. 2022;80:22–32.

18. Newman AAC, Baylis RA, Hess DL, Griffith SD, Shankman LS, Cherepanova OA, Owens GK. Irradiation abolishes smooth muscle investment into vascular lesions in specific vascular beds. JCI Insight. 2018;3.

19. Sörensen I, Adams RH, Gossler A. DLL1-mediated Notch activation regulates endothelial identity in mouse fetal arteries. Blood. 2009;113:5680–5688.

20. He L, Li Y, Li Y, Pu W, Huang X, Tian X, Wang Y, Zhang H, Liu Q, Zhang L, et al. Enhancing the precision of genetic lineage tracing using dual recombinases. Nat. Med. 2017;23:1488–1498.

21. Ikeda J, Scipione CA, Hyduk SJ, Althagafi MG, Atif J, Dick SA, Rajora M, Jang E, Emoto T, Murakami J, et al. Radiation Impacts Early Atherosclerosis by Suppressing Intimal LDL Accumulation. Circ. Res. 2021;128:530–543.

22. Schiller NK, Kubo N, Boisvert WA, Curtiss LK. Effect of γ-Irradiation and Bone Marrow Transplantation on Atherosclerosis in LDL Receptor-Deficient Mice. Arterioscler. Thromb. Vasc. Biol. 2001;21:1674–1680.

23. Zheng X, Liu Z, Bin Y, Wang J, Rao X, Wu G, Dong X, Tong F. Ionizing radiation induces vascular smooth muscle cell senescence through activating NF-κB/CTCF/p16 pathway. Biochim. Biophys. Acta BBA - Mol. Basis Dis. 2024;1870:166994.

24. Mohri Y, Nie J, Morinaga H, Kato T, Aoto T, Yamanashi T, Nanba D, Matsumura H, Kirino S, Kobiyama K, et al. Antagonistic stem cell fates under stress govern decisions between hair greying and melanoma. Nat. Cell Biol. 2025;27:1647–1659.

25. Moradinasab N, Deaton RA, Shankman LS, Owens GK, Brown DE. Label-Efficient Contrastive Learning-Based Model for Nuclei Detection and Classification in 3D Cardiovascular Immunofluorescent Images. In: Xue Z, Antani S, Zamzmi G, Yang F, Rajaraman S, Huang SX, Linguraru MG, Liang Z, editors. Medical Image Learning with Limited and Noisy Data. Cham: Springer Nature Switzerland; 2023. p. 24–34.

