## Supplemental Material for "Dual Lineage Tracing Identifies Cellular Mechanisms Underlying Radiation-Associated Changes in Atherosclerotic Lesion Composition"

**Title**

**Short Title**: Radiation Alters Atherosclerotic Lesion Composition

*Corresponding authors:

**Dr. Gary K. Owens**

University of Virginia School of Medicine

Robert M. Berne Cardiovascular Research Center

PO Box 801394

MR5 Building

Charlottesville, Virginia 22908-1394

**Dr. Rebecca A. Deaton**

University of Virginia School of Medicine

Robert M. Berne Cardiovascular Research Center

PO Box 801394

MR5 Building

Charlottesville, Virginia 22908-1394

**Contents:**

Supplemental Methods
Figures S1-S9
References 5-8

**Supplemental Methods:**

***Validation of bone marrow reconstitution***

Whole blood was collected from control or irradiated mice to assess bone marrow engraftment. Control blood from CD45.1 and *Apoe-/-* (CD45.2) mice were used for single-stain and fluorescence-minus-one controls. Erythrocytes were lysed using BD PharmLyse Lysing Buffer (BD Biosciences, #555899) for 10 minutes at room temperature (RT), followed by centrifugation at 600g for 10 minutes. The lysis step was repeated if visible erythrocyte contamination persisted. Non-specific binding to Fc receptors was blocked using TruStain FcX PLUS (anti-mouse CD16/32) (Biolegend #156604, 1:100) for at least 10 minutes prior to staining. Samples were stained with the following antibodies in phosphate buffered saline (PBS): CD45.1 (BB700, BD Biosciences # 566496, 1:100); CD45.2 (BV605, BD Biosciences #747743, 1:100); and Fixable Viability Dye (eFluor 780, eBiosciences #65-0865-14, 1:1000) for 20-30 minutes at RT in the dark. Following staining, samples were washed at 600g for 5 minutes and resuspended in FACS buffer (1% BSA in PBS). Samples were run on an Attune CytPix cytometer and analyzed using FCS Express v7 with compensation calculated from single-stain controls and Fluorescence Minus One (FMO) controls used for gating.

***Tissue Collection***

After euthanasia by $CO_2$ asphyxiation, mice were gravity-perfused via the left ventricle of the heart with 5 mL of PBS, 10 mL of 4% paraformaldehyde (PFA) (EMS; Catalog# 15710) in PBS, and an additional 5 mL of PBS. Organs were collected and fixed for ≤24 hours in 4% PFA in PBS with gentle shaking. Tissues were transferred to 30% sucrose for up to 48 hours prior to frozen embedding (Epredia Neg-50™, # 6502B). Frozen blocks were stored at -80°C prior to cryosectioning.

***Histology (sectioning, IF, PSR, MOVAT)***

PFA-fixed, frozen embedded tissues were cut in serial 10μm sections. BCA samples were sectioned from the aortic arch to the subclavian bifurcation as previously described. For Immunofluorescence staining, sections were fixed in acetone, air dried, and washed twice in PBS prior to staining for ACTA2 (Alpha-Smooth Muscle Actin Monoclonal Antibody (1A4), Alexa Fluor 405 Novus Biologicals # NBP2-34522AF405; 1:100) and nuclei (DAPI, Invitrogen D21490; 1:100 or NucSpot750, Biotium #41038-T;1:500) and mounting (ProLong™ Diamond Antifade Mountant Invitrogen # P36970). For staining of isolated retina, samples were incubated in DAPI for 10 minutes, washed with PBS, cut for whole mount and mounted as described. Images were acquired in the Advanced Microscopy Facility at the University of Virginia using either a Zeiss LSM880 AiryScan or Leica STELLARIS-8 tauSTED at 20x magnification and 0.6x zoom to acquire a series of z-stack images at 1μm intervals. Acquisition settings were determined using IgG isotype control and were kept constant across images acquired. Maximum intensity projection was used to generate the representative images included in the figures, and Adobe Photoshop was used to process and format those images. Modifications were applied identically across all images. Movat and Picrosirius Red (PSR) staining were performed as previously described.[5–8] Samples were imaged using a Leica Thunder Imager and analyzed using ImageJ as previously described.[5–8] Percent occlusion was determined from Movat stained BCAs images using the following formula: IEL area/lesion area × 100.

***Flow Cytometry Analysis of aortic cells***

Mice were euthanized by $CO_2$ asphyxiation and perfused with 10 mL PBS via the left ventricle of the heart. Whole aortas (including arch, thoracic and abdominal) were excised and placed into FACS buffer (1% BSA in PBS). Each tissue sample was chopped with scissors and digested in 1mL of digestion buffer containing 4 units/ml LiberaseTM (Roche, #355374) and 0.744 units/ml Elastase (Worthington Bio. Corp. LS002279) in RPMI for 60 min at 37°C. Digested samples were collected in 15 ml conical tubes and FACS Buffer was added making a total volume of 13 ml and tubes centrifuged at 1000g for 10 min. The supernatant was discarded and red blood cells in the pellet were lysed using 1ml 1X BD Pharm Lyse Buffer (BD Biosciences Cat no: 555899) for a maximum of 5 minutes. Pellets were washed with FACS buffer and subsequently filtered through 70 µm filters. Remaining cells were stained with viability dye (Invitrogen, eBioscience™ Fixable Viability Dye eFluor™ 780, Cat no: 65-0865-14, 1:1000 dilution) for 20 minutes and fixed with 4% paraformaldehyde for 20 minutes. Samples were run on a Cyteck® Northern Lights™ cytometer and analyzed using FCS Express v7 with compensation calculated from single-stain controls and FMO controls used for gating.

***Cell Profiling analysis (ImageJ)***

Analysis of cellular compositions (*Myh11*-tdTomato$^+$, *Cdh5*-zsGreen$^+$, ACTA2$^+$, *Myh11*-tdTomato$^+$ACTA2$^+$, *Cdh5*-zsGreen$^+$ACTA2$^+$) of healthy BCAs and atherosclerotic lesions was performed using Fiji (ImageJ) software (USA National Institute of Health) and quantified as percent population of either DAPI, Nuclei or ACTA2$^+$ as previously described.[5–8] For validation studies, all vessels contained a minimum of 48 luminal cells and 157 medial cells per vessel. In cases where no reporter-positive cells were detected, limits of detection were estimated using the rule-of-three (3/n), yielding conservative per-vessel detection limits of ~6.3% for luminal tdTomato labeling and ~1.9% for medial zsGreen labeling; study-wide detection limits across SMC-EC dual *Apoe*$^{-/-}$ mice were ~0.38% and ~0.11%, respectively. Post-hoc analysis showed that when stratified by sex, the study was powered to detect differences in reporter expression ≥ 7%.

***scRNAseq***

*Cell processing:* Two samples (control or irradiated) were generated from pooled BCA regions from female SMC-EC dual *Apoe*$^{-/-}$ mice (n=4 for control; n=3 for irradiated) as previously described and summarized below. Mice were euthanized by $CO_2$ asphyxiation and perfused with 20 mL of PBS + 1 µg/mL Actinomycin-D. BCA lesion regions were excised and placed into FACS buffer (1% BSA in PBS) on ice until all samples were collected. Tissues were chopped with scissors and digested in a cocktail containing 4 units/mL Liberase™ (Roche; Catalog# 355374) and 0.744 units/mL Elastase (Worthington Bio. Corp: Catalog# LS002279) in RPMI + 1 µg/mL Actinomycin-D for 60 min at 37°C. Individual samples were pooled and stained for viability (Invitrogen™ eBioscience™ Fixable Viability Dye eFluor™ 780 Cat# 65-0865-14). Live cells were sorted (BD Influx™ Cell Sorter), counted (DeNovix Celldrop™) and submitted directly for scRNAseq library preparation.

*Sequencing, read alignments, and quality control:* All libraries were prepared using 10x Chromium Next GEM Single Cell 3' Kit v3.1 at the Genome Analysis and Technology Core at the University of Virginia, in accordance with the manufacturer's instructions. ~100,000 reads/cell and ~30,000 cells per sample were targeted for a total of 6B reads. Sequencing was performed on the NovaSeq X Plus (Illumina) at Novogene (Sacramento, California) with system specifications of paired-end reads at 150bp length (PE150) and a total data output of 1,800 Gb.

FASTQ files were processed using the Cell Ranger pipeline v9.0.1 (10x Genomics). Resulting reads were aligned to a custom *Mus musculus* GRCm39 reference genome, generated by combining the standard 10x Genomics mouse reference with tdTomato and ZsGreen reporter gene sequences to aid in the detection of SMC- and EC- lineage reporters. Alignment to the reference genome and quantification generated individual filtered gene–barcode matrices for each sample, which were subsequently imported into R (version 4.4.3, 2025-02-28, ucrt) for downstream analysis using the Seurat package (Seurat v5.3.0) with SeuratObject (v5.0.2) and Bioconductor (v3.20). Cells were retained for analysis only if they checked more than 200 and fewer than 5,000 detected genes and displayed mitochondrial transcript levels below 10%.

*Clustering, annotation and analysis:* Gene counts were normalized using log-normalization in Seurat, and highly variable genes were identified using the variance-stabilizing transformation (VST) method. Principal component analysis (PCA) was performed for dimensionality reduction. The number of principal components retained for downstream analysis was determined by inspection of the elbow plot, identifying the point at which additional components contributed minimal additional variance. Based on this assessment, the first 15 principal components, accounting for 83.8% of the total variance, were selected for clustering. Cells were clustered using a graph-based Louvain algorithm, with clustering resolution empirically optimized based on cluster separation and biological interpretability. A resolution of 0.3 was selected to avoid over-partitioning into biologically redundant clusters. Uniform Manifold Approximation and Projection (UMAP) was used for low-dimensional visualization. Feature-level and cluster-level gene expression patterns were visualized using different Seurat functions, including FeaturePlot and DotPlot, with additional figure modification using ggplot2.

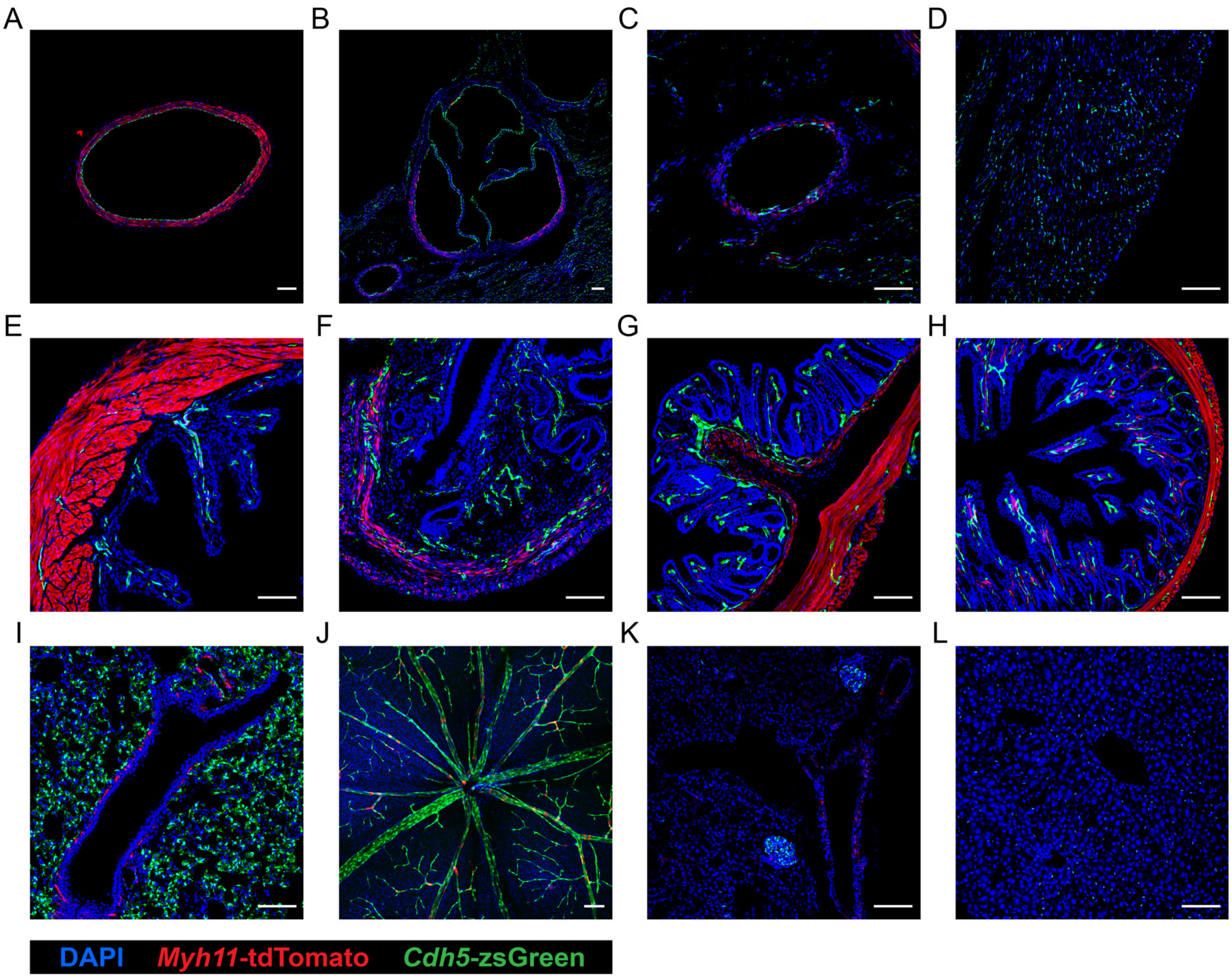


**Figure S1: Cell-type-specific reporter labeling of vascular and visceral smooth muscle cells and endothelial cells across multiple tissues in SMC-EC dual *Apoe*$^{-/-}$ mice.** Tissues were collected from tamoxifen-fed 10-week-old SMC-EC dual *Apoe*$^{-/-}$ mice and analyzed by fluorescence microscopy. Representative images show DAPI$^{+}$ nuclei (blue), tdTomato$^{+}$ smooth muscle cells (red), and zsGreen$^{+}$ endothelial cells (green) in sections of the (**A**) aorta, (**B**) aortic root, (**C**) coronary artery, (**D**) myocardium, (**E**) bladder, (**F**) uterus (females), (**G**) intestine, (**H**) colon, (**I**) lung, (**J**) retina, (**K**) kidney, and (**L**) liver. Scale bar = 100 µm.

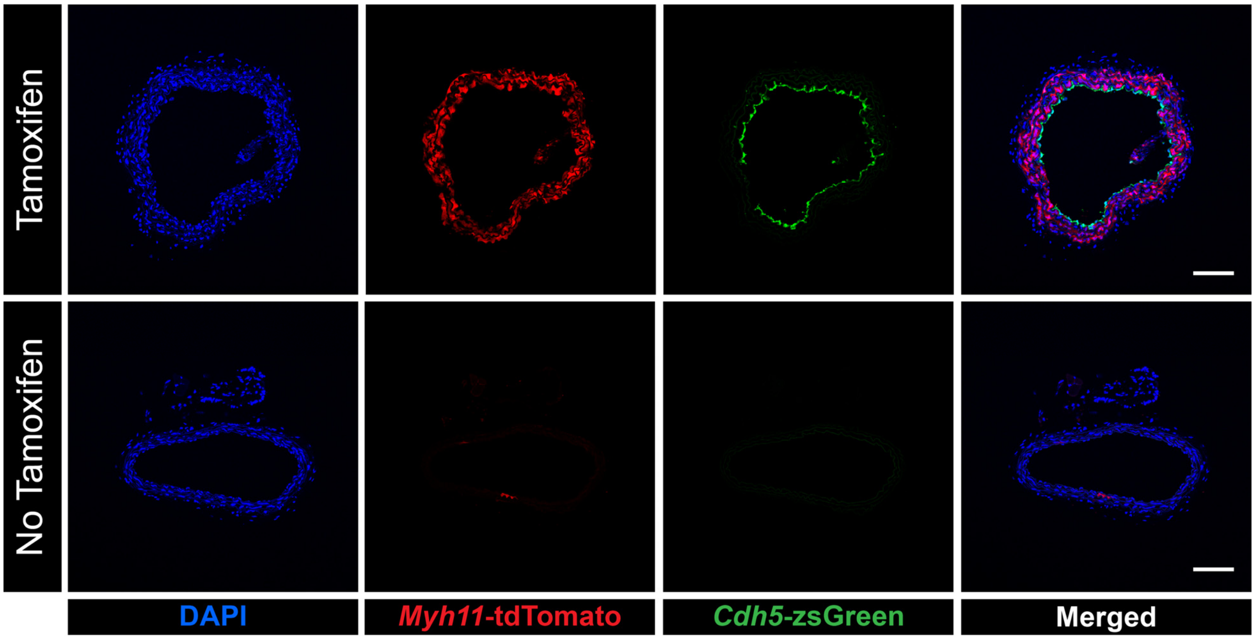


**Figure S2: Tamoxifen-inducible reporter labeling of smooth muscle and endothelial cells in SMC-EC dual *Apoe*$^{-/-}$ mice.** Representative fluorescence images show DAPI$^{+}$ nuclei (blue), tdTomato$^{+}$ smooth muscle cells (red), and zsGreen$^{+}$ endothelial cells (green) in brachiocephalic artery (BCA) sections. Top panels show sections from 10-week-old SMC-EC dual *Apoe*$^{-/-}$ mice fed tamoxifen-containing diet starting at 6-8 weeks of age for 12 days whereas bottom panels show sections from aged-matched mice fed a standard diet. Scale bar = 100 µm.

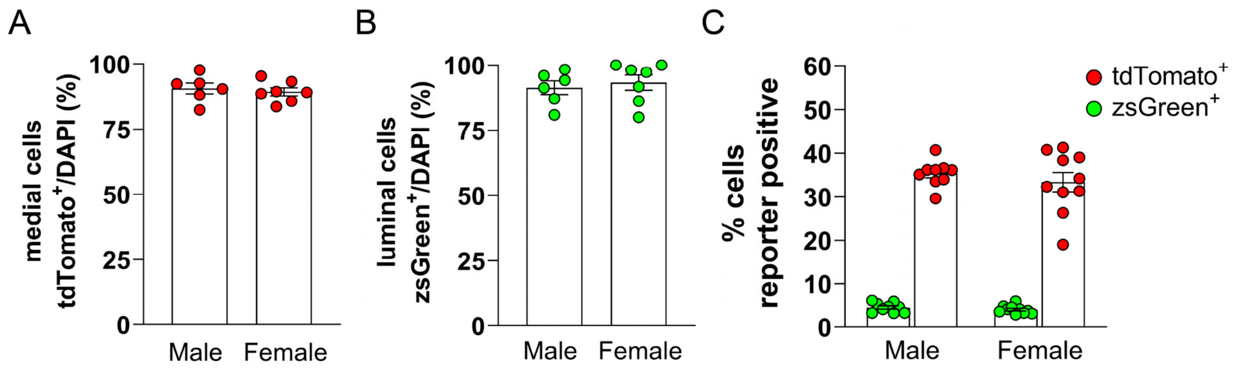


**Figure S3: No sex-dependent differences were detected in smooth muscle or endothelial cell labeling of the brachiocephalic artery from SMC-EC dual *Apoe*$^{-/-}$.** Labeling of cells in the brachiocephalic artery with tdTomato (red circles) or zsGreen (green circles) was quantified following stratification by sex. (**A**) Quantification of the percent tdTomato$^{+}$ medial smooth muscle cells in male versus female mice (P = 0.6202). (**B**) Quantification of the percent zsGreen$^{+}$ luminal endothelial cells in male versus female mice (P = 0.6242). (**C**) Quantification of the percent of aortic cells that are tdTomato$^{+}$ (red circles) or zsGreen$^{+}$ (green circles) in male versus female mice (tdtomato$^{+}$, P = 0.4875; zsGreen$^{+}$, P = 0.9572). Bar graphs show the mean ± SEM, with individual dots representing biologically independent animals. Sample sizes were n=6 males and n=7 females (**A, B**) and n=9 males and n=10 females (**C**). Statistical significance was assessed using an unpaired t-test (**A, B**) or two-way repeated measures ANOVA with Šidák correction for multiple comparisons (**C**).

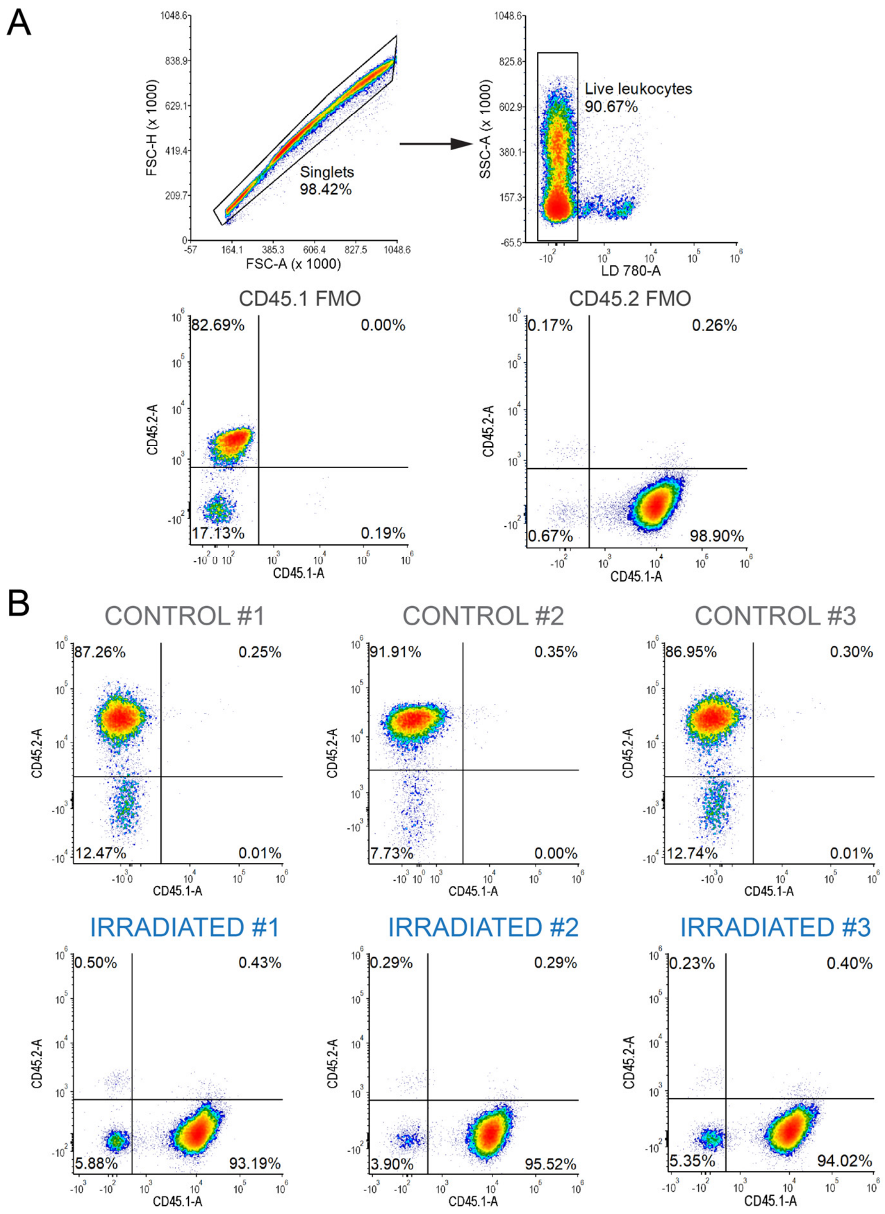


**Figure S4: Validation of bone marrow engraftment following irradiation and BMT.** (**A**) Gating strategy to assess bone marrow engraftment efficiency in peripheral blood leukocytes. Leukocytes were identified by forward and side scatter (FSC/SSC) profiles followed by exclusion of cell doublets and dead cells. Gates for CD45.1 and CD45.2 were set using fluorescence minus one (FMO) controls. (**B**) Representative density plots showing CD45.1 and CD45.2 expression in peripheral blood from three control (top panels) and three irradiated (bottom panels) mice.

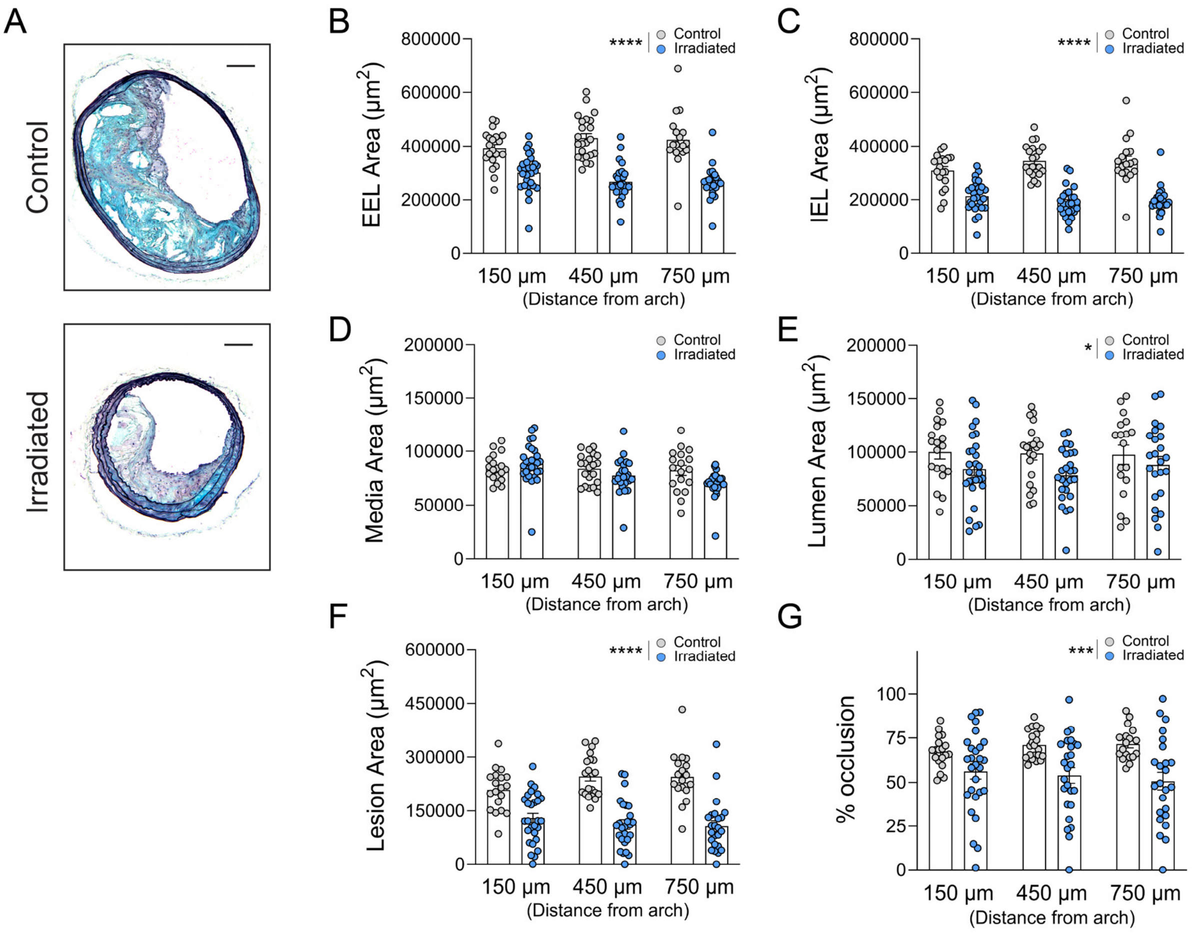


**Figure S5: Irradiation alters vessel remodeling and cross-sectional lesion burden in Western diet-fed control and irradiated mice.** Brachiocephalic artery (BCA) sections from Western diet-fed control (gray circles) and irradiated (blue circles) SMC-EC dual *Apoe*$^{-/-}$ mice were Movat-stained and analyzed at three locations distal to the aortic arch. (**A**) Representative Movat-stained BCA sections collected 450 μm distal to the aortic arch from control (top) and irradiated (bottom) mice. (**B**) Quantification of the external elastic lamina (EEL) area (****P < 0.0001). (**C**) Quantification of the internal elastic lamina (IEL) area (****P < 0.0001). (**D**) Quantification of the media area (P = 0.2644). (**E**) Quantification of the lumen area (*P = 0.0244). (**F**) Quantification of the lesion area (****P < 0.0001). (**G**) Quantification of the percent occlusion (***P = 0.009).

Scale bar = 100 µm (**A**). Bar graphs show the mean ± SEM, with individual dots representing biologically independent animals. Sample sizes vary by location due to occasional tissue loss during histological processing; n values are indicated in the graphs. Statistical significance was assessed using a mixed-effects model with Šidák correction for multiple comparisons, and P values represent the overall treatment effect. (**B-G**).

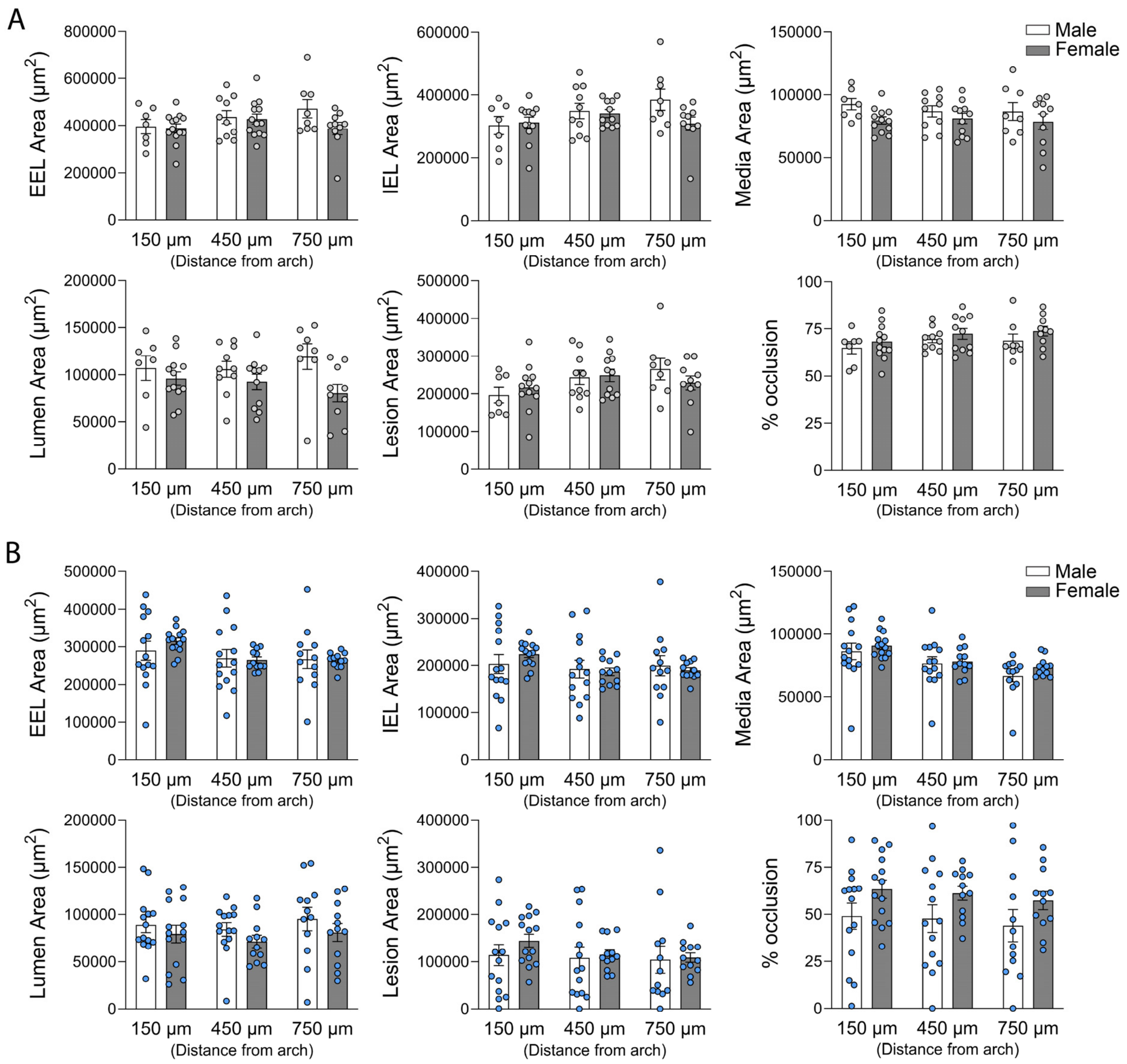


**Figure S6: No sex-dependent differences were detected in vessel remodeling or cross-sectional lesion burden in Western diet-fed control and irradiated mice.** Vessel areas were quantified from Movat-stained brachiocephalic artery sections from Western diet-fed control (gray circles) and irradiated (blue circles) SMC-EC dual *Apoe*$^{-/-}$ mice. (**A**) In control mice, no differences were detected between male and female mice for external elastic lamina (EEL) area (P = 0.1790), internal elastic lamina (IEL) area, (P = 0.179), medial area (P = 0.1756), lumen area (P = 0.0876), lesion area (P = 0.9181), or percent occlusion (P = 0.2690). (**B**) Similarly, no sex-dependent differences were detected in irradiated mice for EEL area (P = 0.7059), IEL area (P = 0.8398), medial area (P = 0.3638), lumen area (P = 0.2023), lesion area (P = 0.5024), or percent occlusion (P = 0.0648). Bar graphs show the mean ± SEM, with individual dots representing biologically independent animals. Sample sizes varied by vessel location due to tissue loss during histological processing; n values are indicated in the graphs. Statistical significance was assessed using a mixed-effects model with Šidák correction for multiple comparisons, and P values represent the overall treatment effect.

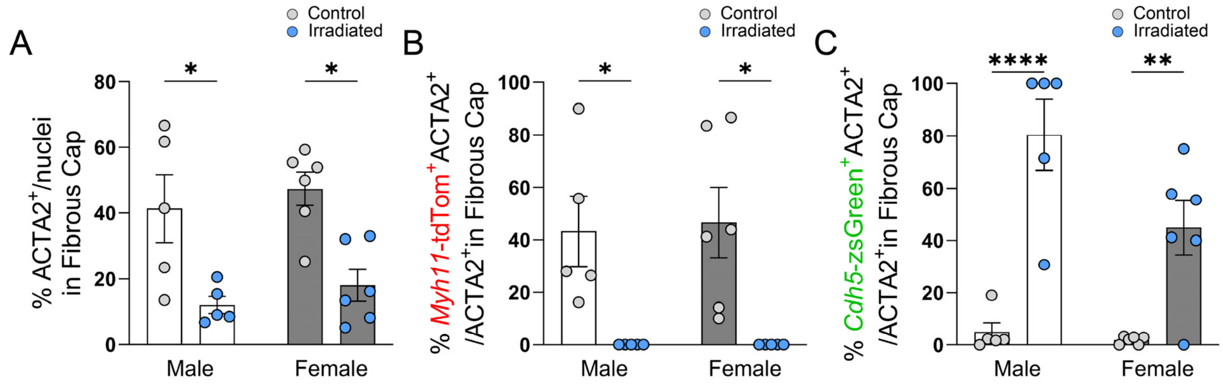


**Figure S7: Sex-stratified analysis of lesion cell composition reveals no sex-dependent differences while preserving treatment effects.** Cell composition of brachiocephalic artery lesions from Western diet-fed control (gray circles) and irradiated (blue circles) SMC-EC dual *Apoe*$^{-/-}$ mice was quantified and analyzed following stratification by sex. (**A**) Quantification of the percent of ACTA2$^{+}$ nuclei relative to total DAPI$^{+}$ nuclei (male, *P = 0.0295; female *P = 0.0147) (**B**) Quantification of tdTomato$^{+}$ ACTA2$^{+}$ cells expressed as a fraction of total ACTA2$^{+}$ cells (male, *P = 0.0492; female *P = 0.0224). (**C**) Quantification of zsGreen$^{+}$ ACTA2$^{+}$ cells expressed as a fraction of total ACTA2$^{+}$ cells (male, ****P < 0.0001; female **P = 0.0091). No significant differences were detected between male (white bars) and female (gray bars) mice within each treatment group following sex stratification. Bar graphs show mean ± SEM, with individual dots representing biologically independent animals. Sample sizes were n = 5 control males, n = 5 irradiated males, n = 6 control females and n = 6 irradiated females (**A-C**). Statistical significance was assessed using a two-way ANOVA with Šidák correction for multiple comparisons.

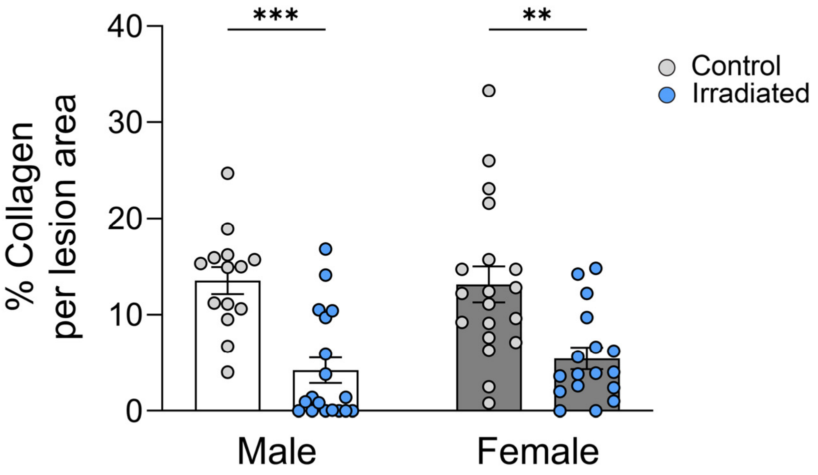


**Figure S8: Sex-stratified analysis of lesion collagen content reveals no sex-dependent differences while preserving treatment effects.** Quantification of the percent collagen content stratified by sex in Western Diet-fed control (gray circles) and irradiated (blue circles) SMC-EC dual *Apoe*$^{-/-}$ mice. No significant differences were detected between male (white bars) and female (gray bars) mice within each treatment group following sex stratification, whereas treatment-dependent differences remained significant within both sexes (male, ***P = 0.0004 and female **P = 0.0023). Bar graphs show mean ± SEM, with individual dots representing biologically independent animals. Sample sizes were n = 14 control males, n = 18 irradiated males, n = 19 control females and n = 17 irradiated females. Statistical significance was assessed using a two-way ANOVA with Šidák correction for multiple comparisons.

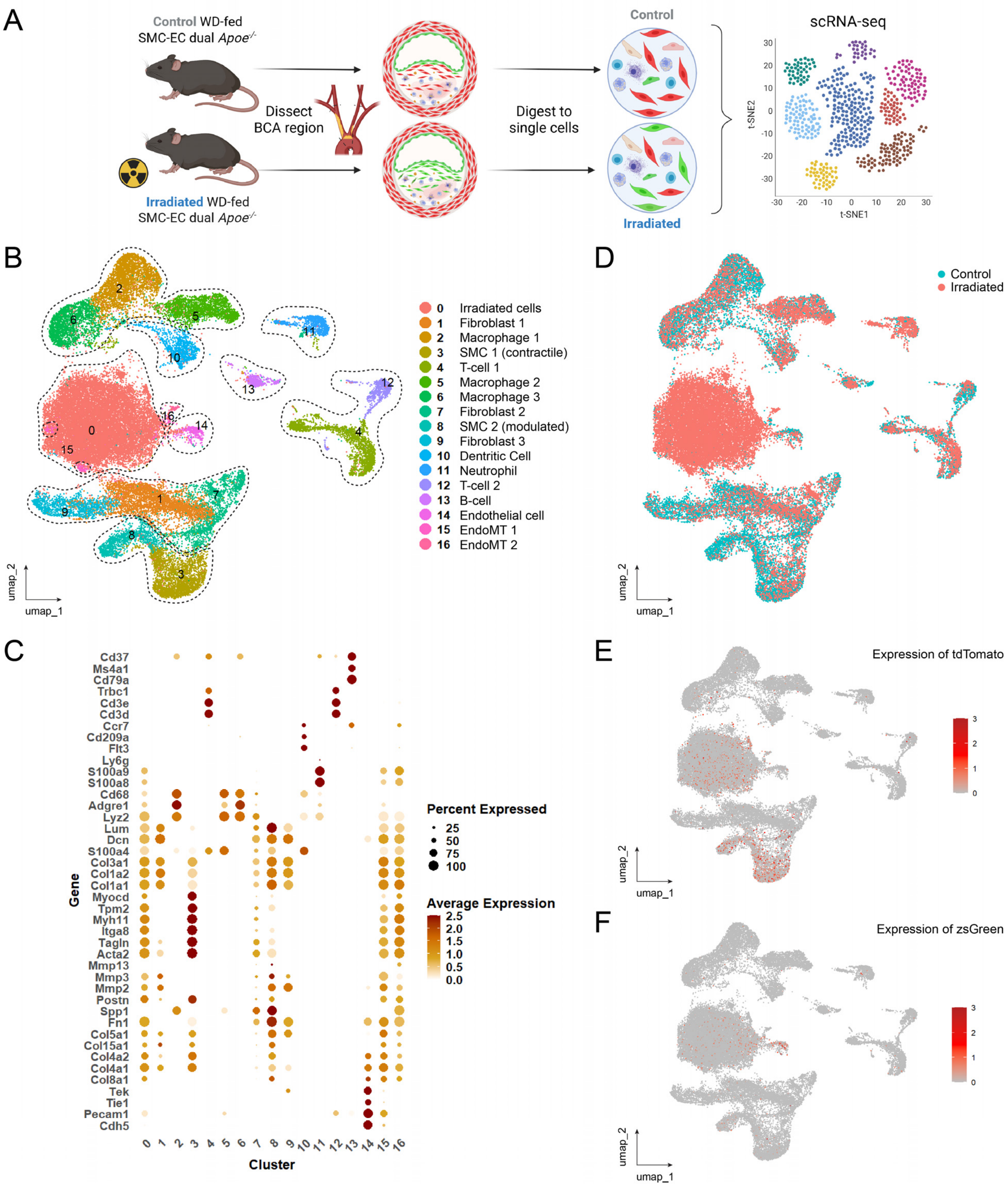


**Figure S9: Single-cell RNA sequencing identified a distinct cell cluster composed almost exclusively of irradiated cells containing lineage-traced smooth muscle and endothelial cells.** Brachiocephalic artery (BCA) regions were isolated from Western-diet fed control (n = 4) and irradiated (n = 3) SMC-EC dual *Apoe*$^{-/-}$ mice. Vessels were enzymatically digested to generate single-cell suspensions, pooled by treatment, and sorted for live cells prior to single-cell RNA sequencing (scRNA-seq) analysis. (**A**) Schematic showing experimental design. (**B**) Uniform Manifold Approximation and Projection (UMAP) visualization of all sequenced cells clustered by cell-type (clusters 0-16) based on canonical marker gene expression. (**C**) Dot plot showing representative marker genes (y-axis) used to define cell clusters (x-axis). Dot size represents the percentage of cells expressing each gene and color intensity indicates average scaled expression. (**D**) UMAP visualization of all cells colored by treatment group (control vs. irradiated) revealing a cluster predominantly composed of cells from irradiated mice (cluster 0). (**E**) UMAP visualization showing expression of tdTomato transcript across all cells. (**F**) UMAP visualization showing expression of zsGreen transcript across all cells.

References:


5. Shankman LS, Gomez D, Cherepanova OA, Salmon M, Alencar GF, Haskins RM, Swiatlowska P, Newman AAC, Greene ES, Straub AC, et al. KLF4-dependent phenotypic modulation of smooth muscle cells has a key role in atherosclerotic plaque pathogenesis. *Nat. Med.* 2015;21:628–637.

6. Newman AA, Serbulea V, Baylis RA, Shankman LS, Bradley X, Alencar GF, Owsiany K, Deaton RA, Karnewar S, Shamsuzzaman S, et al. Multiple cell types contribute to the atherosclerotic lesion fibrous cap by PDGFRβ and bioenergetic mechanisms. *Nat. Metab.* 2021;3:166–181.

7. Cherepanova OA, Gomez D, Shankman LS, Swiatlowska P, Williams J, Sarmento OF, Alencar GF, Hess DL, Bevard MH, Greene ES, et al. Activation of the pluripotency factor OCT4 in smooth muscle cells is atheroprotective. *Nat. Med.* 2016;22:657–665.

8. Gomez D, Baylis RA, Durgin BG, Newman AAC, Alencar GF, Mahan S, St. Hilaire C, Müller W, Waisman A, Francis SE, et al. Interleukin-1β has atheroprotective effects in advanced atherosclerotic lesions of mice. *Nat. Med.* 2018;24:1418–1429.
